# The Desmosome is a Mesoscale Lipid Raft-Like Membrane Domain

**DOI:** 10.1101/401455

**Authors:** Joshua D Lewis, Amber L Caldara, Stephanie E Zimmer, Anna Seybold, Nicole L Strong, Sara N Stahley, Achilleas S Frangakis, Ilya Levental, James K Wahl, Alexa L Mattheyses, Takashi Sasaki, Kazuhiko Nakabayashi, Kenichiro Hata, Yoichi Matsubara, Akemi Ishida-Yamamoto, Masayuki Amagai, Akiharu Kubo, Andrew P Kowalczyk

**Affiliations:** Department of Cell Biology, Emory University School of Medicine, Atlanta, Georgia, USA; Graduate program in Biochemistry, Cell and Developmental Biology, Emory University School of Medicine, Atlanta, Georgia, USA; Graduate program in Cancer Biology, Emory University School of Medicine, Atlanta, Georgia, USA; Buchmann Institute for Molecular Life Sciences, Goethe University Frankfurt, Frankfurt, Germany; Institute for Biophysics, Goethe University Frankfurt, Frankfurt, Germany; Department of Integrative Biology and Pharmacology, University of Texas Health Science Center at Houston, Houston, Texas, USA; Department of Oral Biology, College of Dentistry, University of Nebraska Medical Center, Lincoln, Nebraska, USA; Department of Cell, Developmental, and Integrative Biology, University of Alabama, Birmingham, Alabama, USA; Center for Supercentenarian Medical Research, Keio University School of Medicine, Tokyo, Japan; National Research Institute for Child Health and Development, Tokyo, Japan; Department of Dermatology, Asahikawa Medical University, Asahikawa, Japan; Department of Dermatology, Keio University School of Medicine, Tokyo, Japan; Department of Dermatology, Emory University School of Medicine, Atlanta, Georgia, USA

## Abstract

Desmogleins are cadherin family adhesion molecules essential for epidermal integrity. Previous studies have shown that desmogleins associate with lipid rafts, but the significance of this association was not clear. Here, we report that the desmoglein transmembrane domain (TMD) is the primary determinant of raft association. Further, we identify a novel mutation in the DSG1 TMD (G562R) that causes severe dermatitis, multiple allergies, and metabolic wasting (SAM) syndrome. Molecular modeling predicts that this G to R mutation shortens the DSG1 TMD, and experiments directly demonstrate that this mutation compromises both lipid raft association and desmosome incorporation. Finally, cryo-electron tomography (cryo-ET) indicates that the lipid bilayer within the desmosome is ~10% thicker than adjacent regions of the plasma membrane. These findings suggest that differences in bilayer thickness influence the organization of adhesion molecules within the epithelial plasma membrane, with cadherin TMDs recruited to the desmosome via establishment of a specialized mesoscale lipid raft-like membrane domain.

## Introduction

A characteristic feature of epithelial cells is the assembly of specialized plasma membrane domains that mediate cell adhesion, communication, and barrier function [1, 2]. Among these structures, adherens junctions and desmosomes play overlapping but distinct roles in cell adhesion, signaling, and morphogenesis [1]. Desmosomes are particularly abundant in tissues exposed to mechanical stress, including the skin and heart [3–5]. These adhesive complexes are characterized by highly organized and dense arrangements of desmosomal proteins that can be visualized by electron microscopy [6–8]. Considerable progress has been made in identifying protein interactions that mediate adhesion in both adherens junctions and desmosomes, as well as the associations that anchor these adhesive structures to the cytoskeleton [9–12]. However, the physical constraints imposed by the epithelial plasma membrane that contribute to the segregation of adherens junctions and desmosomal complexes into morphologically, biochemically, and functionally distinct structures are poorly understood.

The adhesive core of the desmosome is comprised of single pass transmembrane desmosomal cadherins termed desmogleins and desmocollins that mediate adhesion between adjacent cells [8, 13, 14]. In humans, there are four desmoglein genes (*DSG1*-*4*), along with three desmocollins (*DSC1*-*3*) [15]. The desmosomal cadherins are coupled to the intermediate filament cytoskeleton through adaptor proteins such as plakoglobin, plakophilins, and the cytolinker protein desmoplakin [6, 16, 17]. These interactions form an electron dense plaque that couples the adhesive interactions of the desmosomal cadherins to the intermediate filament cytoskeleton of adjacent cells, thus conferring tissue resilience to mechanical stress [5, 18]. Loss of desmosome function results in skin [5, 19] and heart [4, 20] diseases characterized by tissue fragility. In the skin, loss of desmosomal adhesion manifests clinically as epidermal blisters and erosions [19, 21], and in some disorders, aberrant thickening of the epidermis [22, 23]. One example of such a disease is severe dermatitis, multiple allergies, and metabolic wasting (SAM) syndrome [24]. This disease is typically caused by null mutations in DSG1, leading to epidermal fragility and barrier defects (Cheng et al., 2016; Has et al., 2015).

We and others have previously demonstrated that desmosomal proteins associate with lipid rafts [25–29]. Lipid rafts are sphingolipid and cholesterol enriched membrane microdomains that introduce spatial heterogeneity into lipid bilayers [30–33]. These domains are critical for protein trafficking, membrane organization, and signaling [34–38]. The sphingolipids present in rafts feature long saturated acyl chains that, along with cholesterol, contribute to the more ordered, densely packed, and thicker membrane environment characteristic of lipid rafts [33, 34, 39]. Desmogleins and other desmosomal proteins have been shown to associate with lipid raft membrane domains as determined by detergent resistance and buoyancy on sucrose gradients [25, 26, 28, 29]. In addition, disruption of lipid rafts by removal of cholesterol from cellular membranes results in weakened desmosomal adhesion, suggesting that lipid rafts play a role in desmosome homeostasis [25, 26]. However, we do not know how desmosomal cadherins target to raft domains or how incorporation into raft domains impacts desmosomal cadherin function.

In the present study, we sought to determine the mechanisms by which raft association governs desmosome assembly, and to identify the determinants of desmoglein partitioning to rafts. Our results indicate that the transmembrane domain (TMD) of the desmogleins is critical for raft association, and that the E-cadherin TMD does not support raft targeting. Raft association appears to be essential for desmoglein function, as a novel mutation that shortens the TMD of human DSG1 abrogates lipid raft targeting, impairs desmosome association, and causes the human skin disease SAM syndrome. Cryo-electron tomography and sub-tomogram averaging demonstrates that the lipid bilayer within the desmosome is thicker than the adjacent plasma membrane, consistent with predictions that the lipid bilayer is thicker at raft domains compared to non-raft membranes. Thus, our results support a model in which the desmosome is a specialized type of lipid raft membrane microdomain, and that the lengthy desmoglein TMD enables efficient desmosome incorporation by facilitating desmoglein partitioning into the thicker desmosomal lipid bilayer. These findings suggest that epithelial junctional complexes achieve plasma membrane domain specification not only through selective protein interactions, but also through constraints imposed by the biophysical characteristics of the plasma membrane.

## Results

### Palmitoylation of Dsg3 is not required for lipid raft association

Desmogleins and other desmosomal components associate with lipid raft membrane microdomains [25–28]. A number of raft associating proteins, including plakophilins, utilize palmitoylation as a membrane raft targeting mechanism [40–42]. Palmitoylation is a reversible post-translational modification that occurs when palmitoyltransferases add a 16-carbon fatty acid (palmitate) to cysteine residues [43, 44]. Sequence alignments (Figure 1A) reveal that desmosomal cadherins contain conserved cysteine residues at the cytoplasmic face of the transmembrane domain, and our previous studies have shown that these residues are critical for desmoglein palmitoylation [45]. We hypothesized that palmitoylation of desmogleins would mediate lipid raft association. Therefore, we mutated cysteines 640 and 642 to alanine residues in murine Dsg3 and used a lentiviral expression system to generate stable A431 cell lines expressing FLAG tagged wild type or mutant Dsg3(CC). Mass tag labeling confirmed that the mutation of these conserved membrane proximal cysteines eliminated Dsg3 palmitoylation (Figure 1B). Interestingly, the loss of Dsg3 palmitoylation had no discernable effect on lipid raft association as determined by Dsg3 incorporation into buoyant and detergent resistant membranes (DRM) [46] (Figure 1C). In addition, both WT Dsg3 and Dsg3(CC) localized to cell-cell borders as assessed by widefield immunofluorescence (Figure 1D). Furthermore, Dsg3 and Dsg3(CC) exhibited similar Triton-X 100 solubility, suggesting no defect in the desmosome or cytoskeletal association of Dsg3(CC) (Figure 1E-1G). Collectively, these results indicate that palmitoylation is not required for lipid raft association of desmogleins or for normal Dsg3 subcellular distribution in quiescent A431 monolayers.

**Figure 1:**
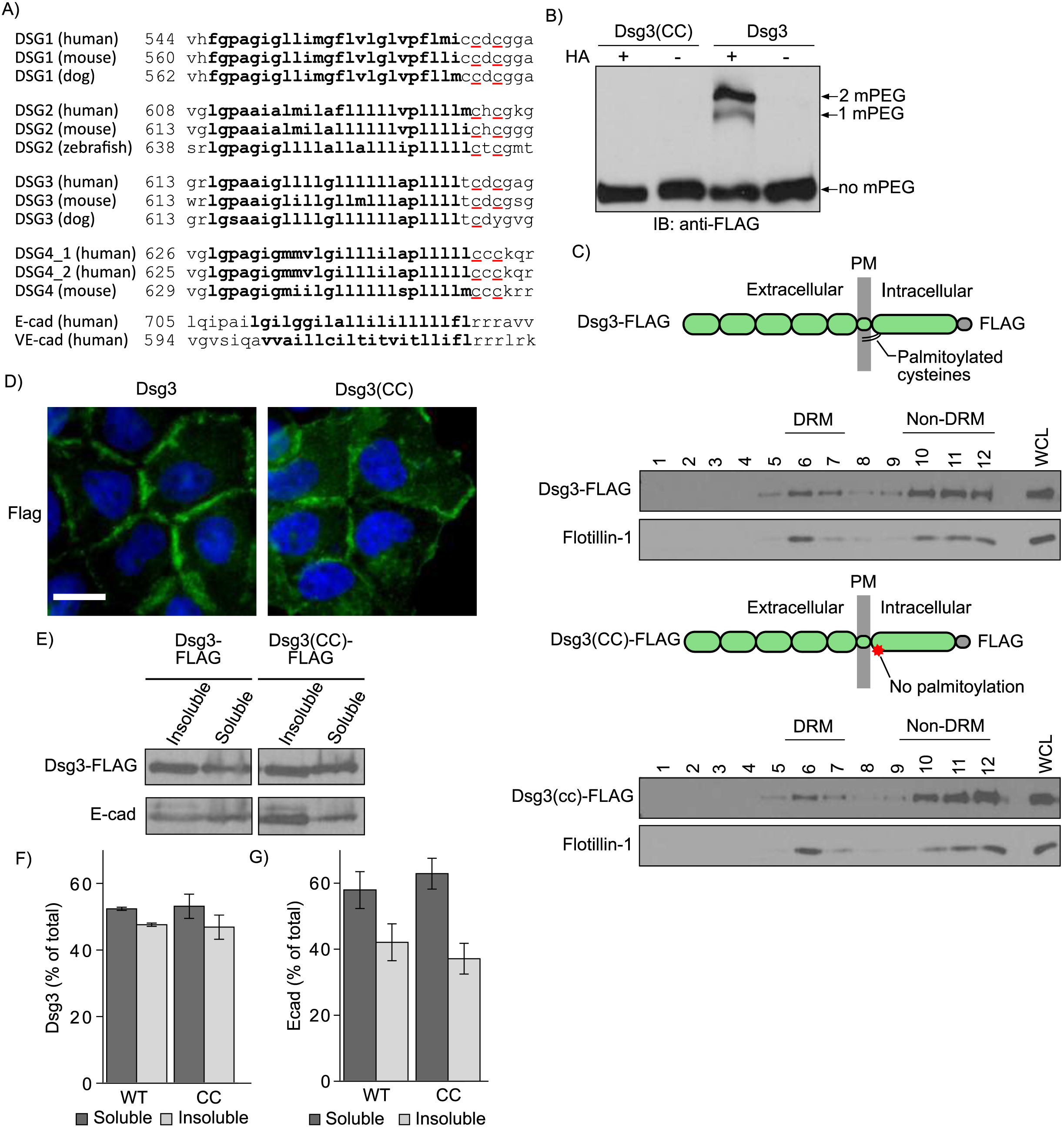
Palmitoylation is not required for Dsg3 lipid raft association. A) Sequence alignment of the desmogleins reveals a pair of highly conserved cysteine residues (yellow) at the interface between the transmembrane domain (light blue) and the cytoplasmic domain. B) Mass-tag labeling replaces palmitoyl moieties on cysteine residues with mPEG, causing a size shift detectable by western blot analysis. Dsg3 is doubly palmitoylated and mutation of the membrane-proximal cysteine residues to alanine abolishes palmitoylation. C) Lipid raft fractionation of HeLa cells expressing Dsg3-FLAG from adenoviruses reveals no defect in lipid raft targeting of the palmitoylation-null mutant. D) Widefield images of A431 cell lines stably harboring flag tagged constructs. Scale bar = 20μm. E) Western blot of Triton-X 100 soluble pools and insoluble pools from A431 cell lines stably expressing either Dsg3 or Dsg3(EcadTMD). F) Densitometry quantification of Dsg3 in triton soluble and insoluble pools from Panel E. Loss of palmitoylation has no detectable effect on the solubility of Dsg3 in Triton X-100, a classic measure of desmosome and cytoskeletal association. F) Densitometry quantification of E-cadherin distribution between Triton soluble and insoluble pools.

### The transmembrane domain of desmogleins mediates lipid raft association

In addition to palmitoylation, emerging evidence indicates that lipid raft association of membrane spanning proteins is also regulated by the physiochemical properties of the transmembrane domain (TMD) [47]. In particular, TMD length is a critical determinant for targeting to lipid rafts [47–50]. Sequence alignments (Figure 1A) indicate that the TMDs of the desmogleins, which associate with rafts, are considerably longer (24 amino acids) than the corresponding TMDs of classical cadherins, such as E-cadherin (21 amino acids) and VE-cadherin (20 amino acids), which exhibit minimal raft association [25]. Recent studies indicate that the free energy of raft association can be calculated based on TMD length, surface area, and palmitoylation [47]. These parameters predict efficient WT Dsg1 raft partitioning (∆G_raft_=0.17), with markedly lower raft affinity for the TMD of E-cadherin (∆G_raft_=0.30) (Table 1). To directly test if the TMD is the principle motif conferring lipid raft association on the desmoglein family of proteins, we generated a chimeric cadherin in which the Dsg3 TMD was replaced with the E-cadherin TMD (Dsg3(EcadTMD)). Lentiviral transduction was used to generate stable A431 cell lines expressing either wild type Dsg3-FLAG or Dsg3(EcadTMD)-FLAG. Sucrose gradient fractionations demonstrated that the Dsg3(EcadTMD) chimera was virtually excluded from DRM fractions when compared to wild type Dsg3 (Figure 2A and 2B). Immunofluorescence localization indicated that Dsg3(EcadTMD) localized to cell-cell contact sites. However, Triton-X 100 extraction showed decreased insoluble pool partitioning as assessed by both immunofluorescence (Figure 2C) and western blot analysis (Figure 2D-2F), suggesting decreased Dsg3(EcadTMD) association with cytoskeletal elements relative to wild type Dsg3. Expression of the Dsg3(EcadTMD) mutant caused no apparent changes in endogenous E-cadherin distribution (Figure 2A, 2D-2F). To determine if the Dsg3 TMD is sufficient to confer lipid raft targeting, we constructed interleukin 2 receptor (IL2R) α chain-Dsg3 chimeric proteins comprising the IL2R extracellular domain coupled to the Dsg3 cytoplasmic tail with either the IL2R TMD or the Dsg3 TMD (Figure 2G and reference 51). The IL2R-Dsg3 chimera harboring the Dsg3 TMD partitioned to DRM fractions, whereas the chimera containing the IL2R TMD did not partition with DRM fractions. Collectively, these studies indicate that the Dsg3 TMD is the primary determinant of Dsg3 raft association.

**Table 1.**
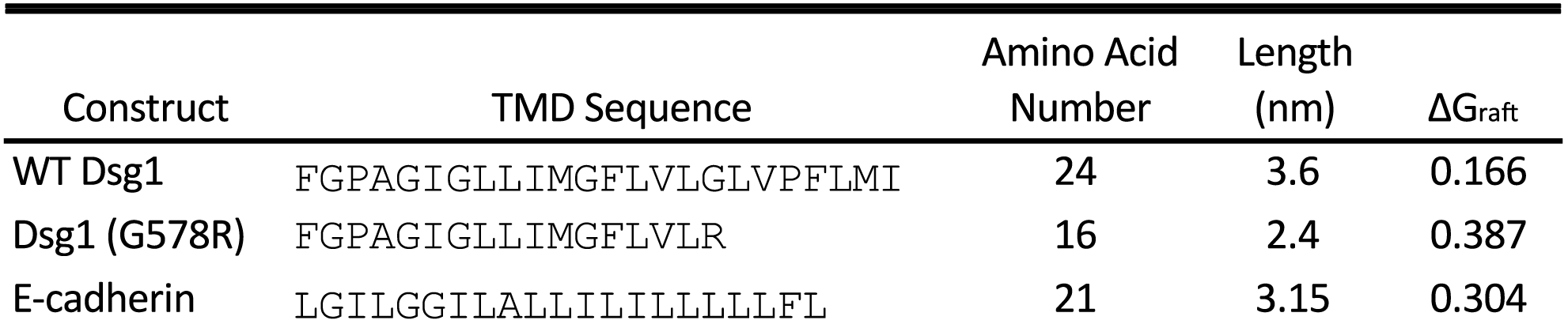
Summary of Transmembrane domains

**Figure 2:**
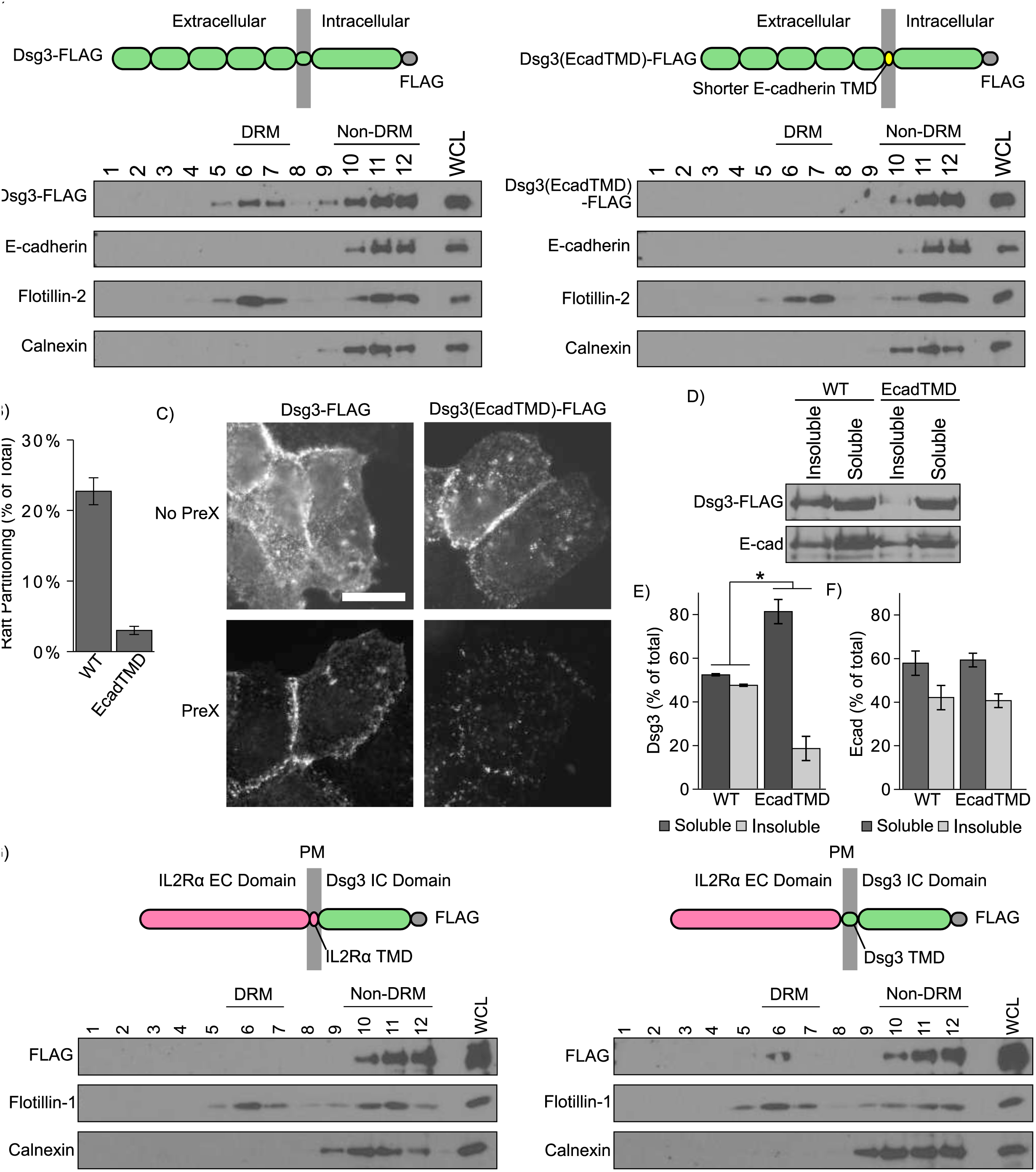
The Dsg3 TMD is necessary for lipid raft association. A) Sucrose gradient fractionation of A431 cells stably expressing murine Dsg3 (wild-type or EcadTMD mutant). Replacing the Dsg3 TMD with the shorter E-cadherin TMD (yellow) abolishes lipid raft targeting. B) Densitometry quantification of WT Dsg3 and Dsg3(EcadTMD) in detergent resistant membrane (DRM) fractions shown in Panel A. C) Dsg3(EcadTMD) is more susceptible than Dsg3 WT to pre-extraction in Triton X-100 prior to fixation and immunofluorescence localization. Scale bar = 20μm. D) Western blot analysis indicates Dsg3(EcadTMD) is more soluble in Triton X-100 than Dsg3 WT. E) Quantification of Dsg3 in Triton soluble or insoluble pools from Panel D. F) Quantification of E-cadherin in Triton soluble and insoluble pools in cells expressing either WT Dsg3 or the Dsg3(EcadTMD) mutant G) Lipid raft fractionation of FLAG-tagged IL2R-Dsg3 chimeras expressed in HeLa cells using an adenoviral delivery system. Inclusion of the lengthy Dsg3 TMD in the chimera (right panel) confers lipid raft targeting. *p<0.05

To test if the TMD of other desmoglein family members also functions in raft association, similar experiments were conducted in the context of Dsg1. Dsg1 WT and a Dsg1(EcadTMD) chimera were generated. Both proteins were tagged with a carboxyl terminal green fluorescent protein (GFP) and stably expressed in A431 cell lines as described above. Similar to the Dsg3(EcadTMD) chimera, the Dsg1(EcadTMD) chimera showed a marked decrease in association with DRM fractions as determined by sucrose gradient fractionation (Figure 3A and 3B). Additionally, Dsg1(EcadTMD) was partially excluded from Triton insoluble fractions of cell lysates (Figure 3C-3E), similar to the results seen with Dsg3(EcadTMD) (Figure 2D-2F). Lastly, both the WT Dsg1 and Dsg1(EcadTMD) demonstrated border staining characteristic of desmogleins (Figure 3F). Together, these results demonstrate a central role for the TMDs of the desmoglein family in lipid raft association.

**Figure 3:**
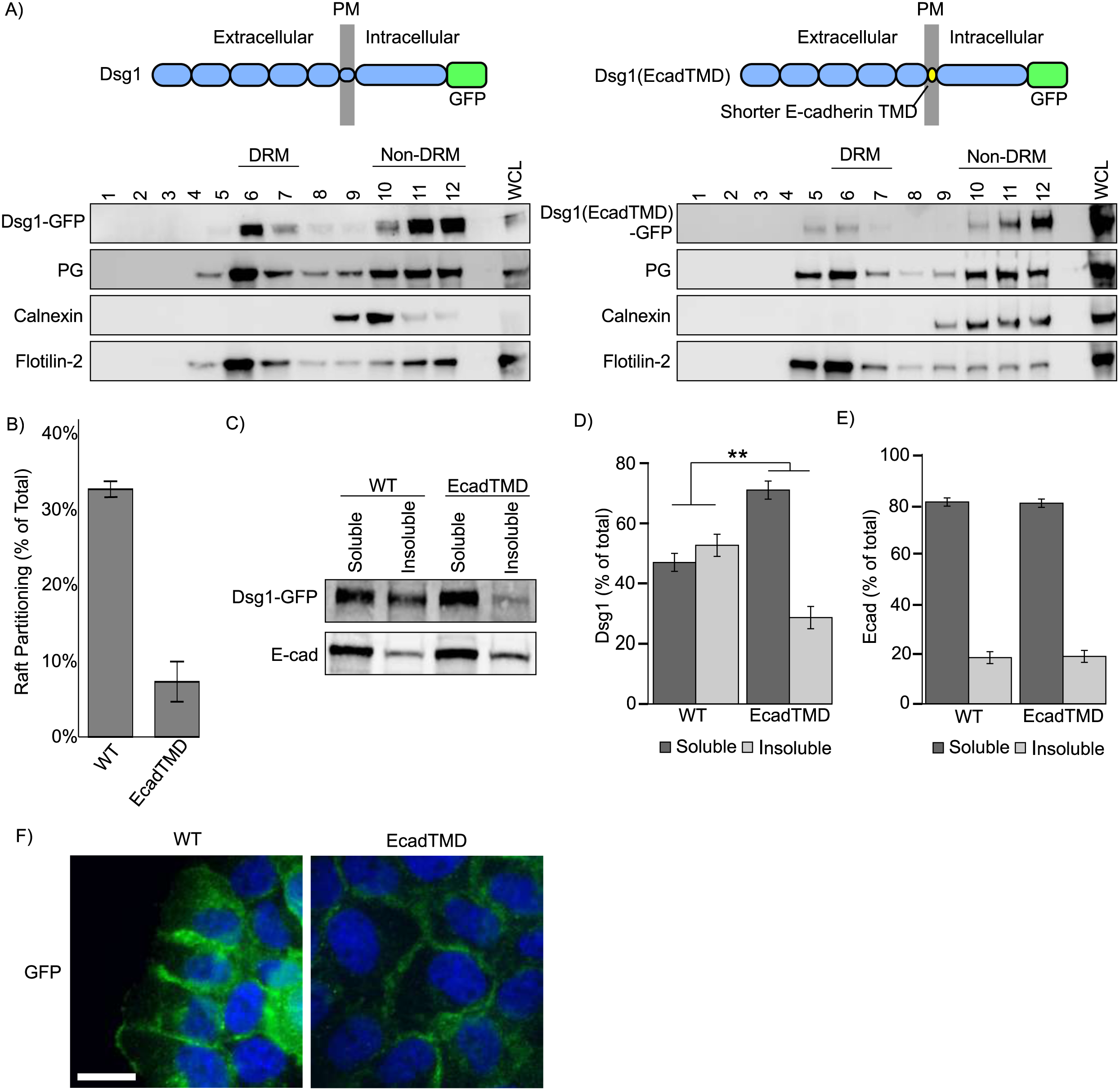
The Dsg1 TMD is critical for lipid raft association. A) Western blot of Triton-X 100 extracts and sucrose gradient fractionations of A431 cells stably expressing murine WT Dsg1 (Dsg1(EcadTMD) chimera. B) Quantification from densitometry analysis of the percentage of total Dsg1 in the detergent resistant membrane (DRM) fractions of sucrose gradient fractionations. C) Differential detergent extraction and western blot analysis indicates that Dsg1(EcadTMD) is more soluble in Triton X-100 than wild type Dsg1. D) Quantification of Dsg1 western blots shown in panel C. E) Quantification of E-cadherin western blots shown in panel C. F) Widefield images of A431 cells expressing either GFP tagged WT Dsg1 or Dsg1(EcadTMD). Scale bar = 20μm. **p<0.001

### A mutation in the transmembrane domain of DSG1 causes severe dermatitis, multiple allergies, and metabolic wasting syndrome

Loss of DSG1 function is associated with a number of autoimmune, infectious, and genetic diseases [19, 20, 22]. One recently discovered desmosome associated disease is severe dermatitis, multiple allergies, and metabolic wasting (SAM) syndrome [24]. Most instances of SAM syndrome are caused by homozygous functional null mutations in the desmosomal cadherin desmoglein 1 (*DSG1*) [52, 53]. Here, we report a novel and dominantly inherited heterozygous DSG1 missense mutation within the DSG1 TMD (Figure 4). The probands presented with ichthyosiform erythrokeratoderma, diffuse palmoplantar keratosis and multiple allergies (Figure 4A). Proband III-2 suffered metabolic wasting and died of status asthmaticus and recurrent infections. Hemotoxylin and eosin staining of skin biopsied from the proband revealed compact hyperkeratosis with parakeratosis, frequent detachment of the entire stratum corneum, and dissociation of individual corneocytes (Figure 4B). Although these findings indicate an adhesion defect, we observed minimal alterations in desmosome ultrastructure when patient epidermis was examined by electron microscopy (Figure 4C). These clinical and genetic observations led us to diagnose the patient with SAM syndrome. Unlike previously reported instances of *DSG1* mutations in SAM syndrome [24, 52–54], this patient harbored a novel missense mutation in DSG1 which introduces a hydrophilic arginine residue (p.G562R) into the otherwise hydrophobic transmembrane domain of DSG1 (Figure 4D-4F). Subsequent to our characterization of this initial family, a second unrelated individual was identified with a G562R heterozygous mutation previously reported as a case of erythrokeratoderma variabilis [55]. The parents of this patient lacked this mutation and were disease free. Together, these observations demonstrate that a heterozygous G562R mutation in the DSG1 TMD causes a human skin disease best characterized clinically as SAM syndrome.

**Figure 4:**
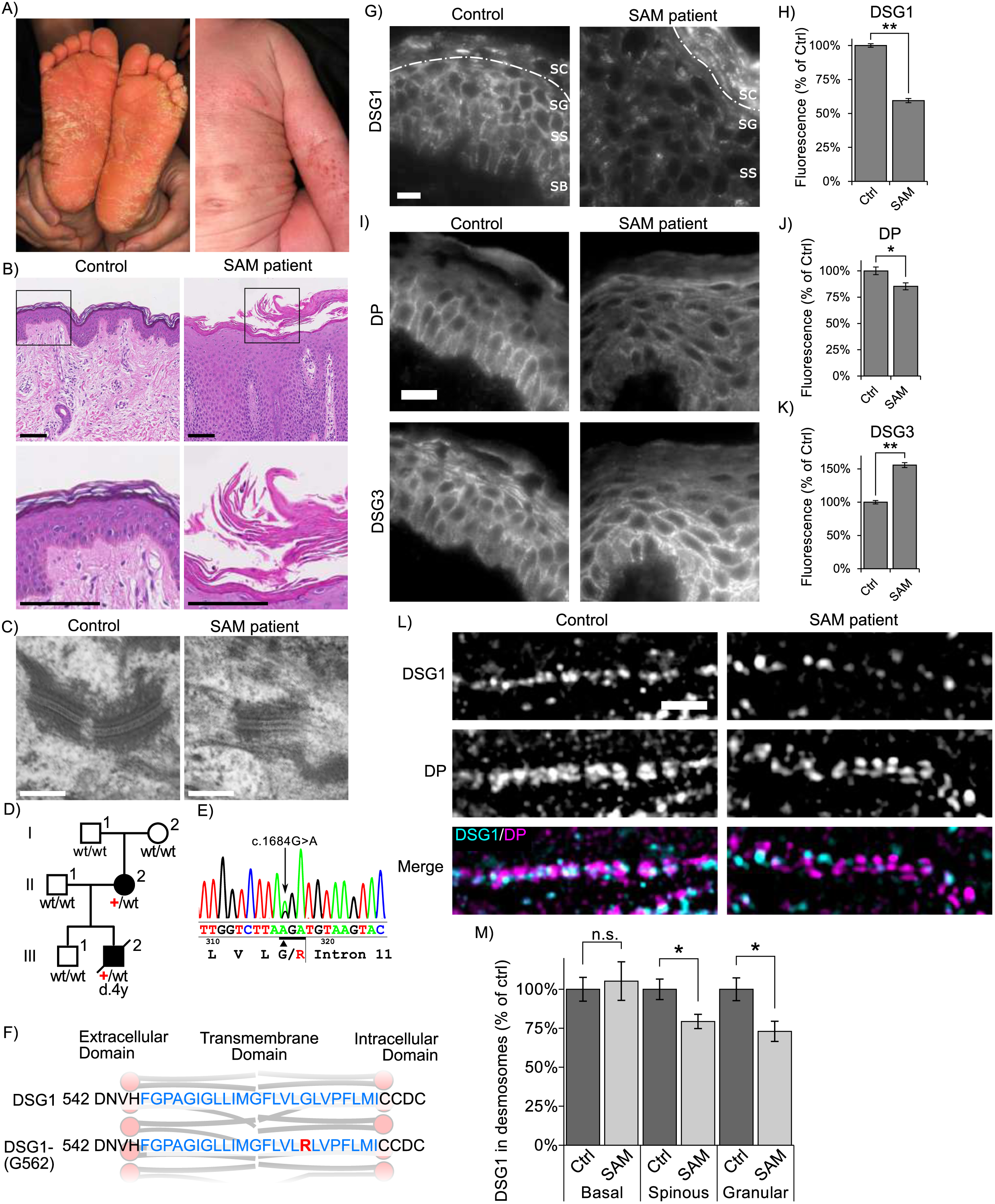
Desmoglein 1 (DSG1) transmembrane domain mutation causes severe dermatitis, multiple allergies, and metabolic wasting (SAM) Syndrome. A) Individual III-2 displays feet covered with hyperkeratotic yellowish papules and plaques, and ichthyosiform erythroderma with severe itch occur over much of his body. B) Hemotoxylin and eosin staining of III-2’s skin biopsy reveals acantholytic lesions in the upper layers of the epidermis. Scale bar = 100 μm. C) Electron micrographs of epidermal sections from the proband indicate relatively normal desmosome morphology. Scale bar = 200 nm. D) Pedigree of affected individuals and near relatives. Inheritance determined by genomic DNA sequencing. E) Genomic DNA sequencing of white blood cells reveals these SAM patients have a heterozygous point mutation, c.1684G>A (black arrow) in DSG1. The adjacent splice site is unaffected. F) Schematic showing the location of the SAM-causing G-to-R substitution (red) within the transmembrane domain (blue). G-H) Widefield microscopy of DSG1 immunofluorescence in human skin biopsies reveals both DSG1 downregulation and inappropriate clustering at cell borders in SAM patient epidermis. SC = stratum corneum, SG = stratum granulosum, SS = stratum spinosum, SB = stratum basale. SC/SG boundary demarcated by dashed line. Downregulation of DSG1 is observed in the SG and SS. I-K) Desmoplakin (DP) is slightly downregulated in patient skin, and DSG3 is upregulated. Scale bar = 20 μm. L-M) Structured Illumination Microscopy (SIM) indicates reduced desmosomal DSG1 in SAM patient tissue in the stratum spinosum and granulosum. Scale bar = 5 μm. *p<0.01, **p<0.001

To determine how the G562R mutation impacted DSG1 organization in patient skin, biopsies from the proband were processed for immunofluorescence microscopy. DSG1 levels were markedly reduced (~40%) in the spinous and granular layers of patient epidermis (Fig 4 G and H), and DSG1 localized in cytoplasmic puncta and aberrant clusters at cell-cell borders. Interestingly, DSG1 staining in patient stratum corneum was markedly increased, perhaps reflecting increased antibody penetration. Desmoplakin levels were slightly reduced in patient epidermis, whereas DSG3 levels were markedly increased (Figure 4I-4K). To further investigate alterations in DSG1 distribution, we performed structured illumination microscopy (SIM) on patient and control epidermis. DSG1 fluorescence intensity within patient and control desmosomes was comparable in basal keratinocytes, where DSG1 expression is low and other DSG isoforms (DSG2, DSG3) are expressed. However, DSG1 fluorescence intensity in patient desmosomes was significantly reduced (Figure 4L and 4M) in the spinous and granular layers where DSG1 is prominently expressed. Thus, although morphologically normal desmosomes could be observed by electron microscopy (Figure 4C), these desmosomes apparently lack sufficient DSG1 levels to support normal epidermal cohesion.

To investigate the mechanism by which the DSG1(G562R) mutation causes SAM syndrome, GFP-tagged murine wild type Dsg1α and a mutant harboring the equivalent G-to-R substitution, Dsg1(G578R), were expressed in A431 epithelial cells. Widefield fluorescence imaging revealed that both WT Dsg1 and Dsg1(G578R) were present at cell-to-cell borders (Fig 5A). Interestingly, the Dsg1(G578R) mutant also exhibited a prominent perinuclear staining pattern. There were no obvious differences in desmoplakin localization in the two cell lines (Figure 5A), and plakoglobin generally co-localized with both cell-cell border and perinuclear pools of WT Dsg1 and Dsg1(G578R) (Figure 5B). To determine if the Dsg1(G578R) mutant was defective in desmosome targeting, SIM was performed and Dsg1 fluorescence intensity was measured at cell borders both within and outside of individual desmosomes to control for possible variations in Dsg1 levels at different cell-cell contact sites. We observed that Dsg1(G578R) displayed a decreased association with desmosomes when compared to WT Dsg1 (Figure 5C and 5D). Furthermore, parallel bands of GFP fluorescence signal overlapping with DP were routinely observed for WT Dsg1-GFP but not for Dsg1(G578R)-GFP (Figure 5C and 5E). Lastly, WT Dsg1 efficiently entered a detergent resistant pool, consistent with incorporation into insoluble desmosome and cytoskeletal-associated complexes, whereas mutated Dsg1(G578R) remained predominantly soluble (Figure 5F-5H). Together, these findings indicate that the G-to-R TMD mutation reduces DSG1 incorporation into desmosomes in both cultured cells and in patient epidermis.

**Figure 5:**
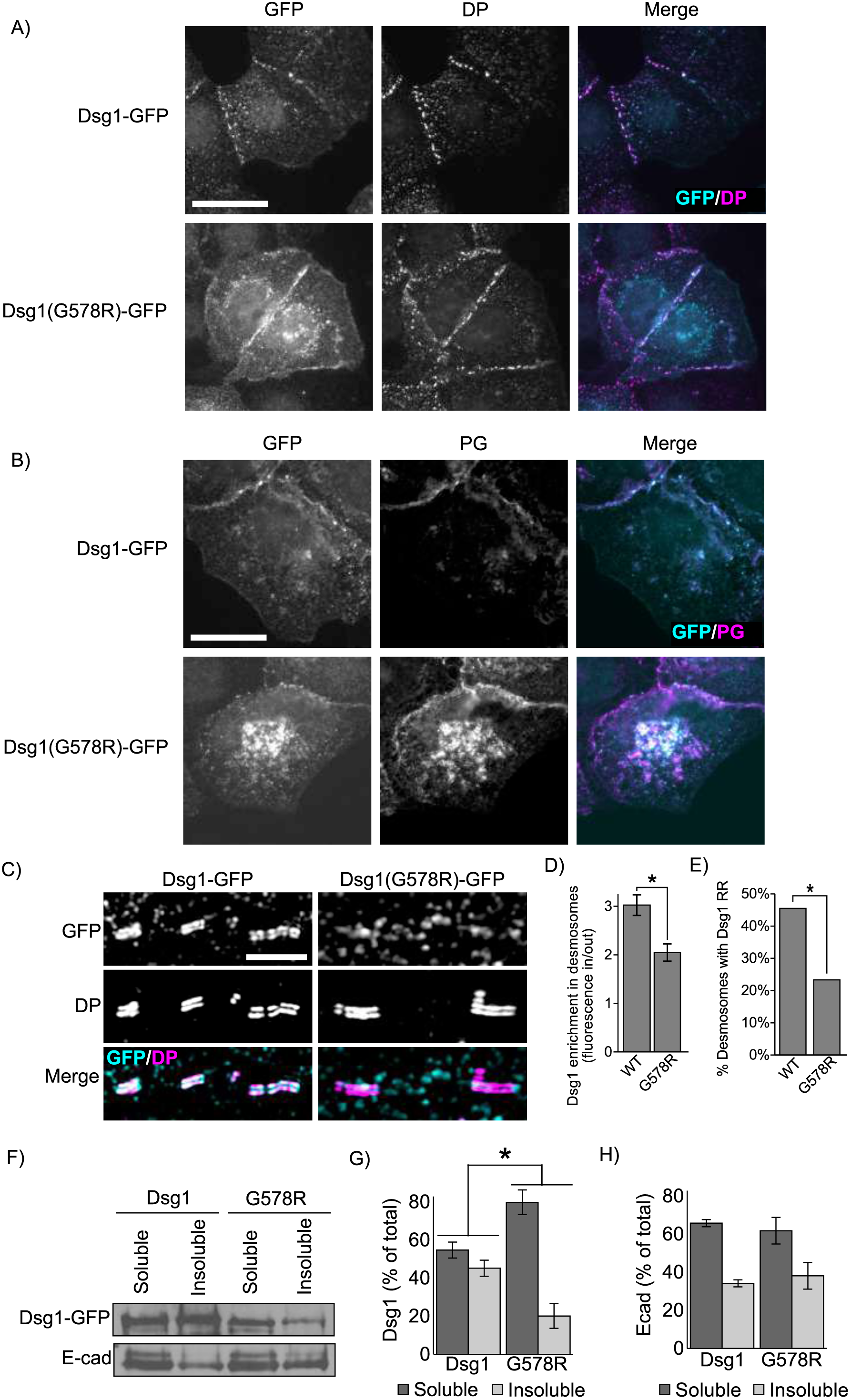
SAM-causing DSG1 mutation causes defects in junction targeting. A) Widefield immunofluorescence micrographs of A431 cell lines stably expressing either wild type Dsg1-GFP or Dsg1(G578R)-GFP reveal broadly similar distribution of desmoplakin (DP) and B) colocalization between DSG1 and plakoglobin (PG). Scale bar = 20 μm. C) Super-resolution micrographs of A431 stable cell lines acquired using structured illumination microscopy (SIM) reveal defects in Dsg1(G578R) desmosome targeting. Scale bar = 5 μm. D) Desmosomes are identified by SIM imaging as regions of parallel desmoplakin staining resembling rail road tracks. Quantification of Dsg1 found within DP railroad tracks compared to border Dsg1 outside of railroad tracks. E) Quantification of railroad track appearance observed for WT or mutant Dsg1.GFP. F-G) The G578R mutation increases solubility of the mutant in Triton X-100 as determined by western blot analysis. E-cadherin distribution in Triton soluble and insoluble pools is not altered in A431 cell lines expressing the Dsg1 mutant. *p<0.05

In addition to being deficient in desmosome targeting, we also observed that DSG1 was present in cytoplasmic puncta in SAM patient epidermis (Figure 4G) and that Dsg1(G578R) was concentrated in perinuclear compartments in A431 cell lines (Figure 5A and 5B). To determine if the Dsg1(G578R) mutant exhibited membrane trafficking defects, cell surface proteins were biotinylated and pulse chase experiments conducted to measure Dsg1 turnover rates. These experiments revealed no difference in the rate of Dsg1(G578R) turnover from the plasma membrane compared to WT Dsg1 (Figure 6A and 6B). To measure rates of delivery to the plasma membrane, cell surface proteins were cleaved using trypsin and the rate of Dsg1 recovery at the cell surface was monitored by biotinylation (Figure 6C and 6D). Prior to trypsinization, Dsg1(G578R) cell surface levels were similar to WT Dsg1(Figure 6E), indicating that steady state cell surface levels of the mutant were comparable to WT Dsg1. However, while the surface pool of WT Dsg1 recovered within 3-6 hours after trypsinization, Dsg1(G578R) exhibited delayed plasma membrane recovery (Figure 6D). To determine if Dsg1(G578R) was being retained in secretory compartments, A431 cell lines were grown in low calcium medium overnight to internalize all cadherins, and subsequently switched to high calcium medium to allow Dsg1 to traffic out to cell-cell borders. These experiments revealed that Dsg1(G578R) was retained in GM130-labeled compartments (Figure 6F and 6G), indicating that the G-to-R mutation causes retention of Dsg1 in the Golgi apparatus, delaying its trafficking through the secretory pathway.

**Figure 6:**
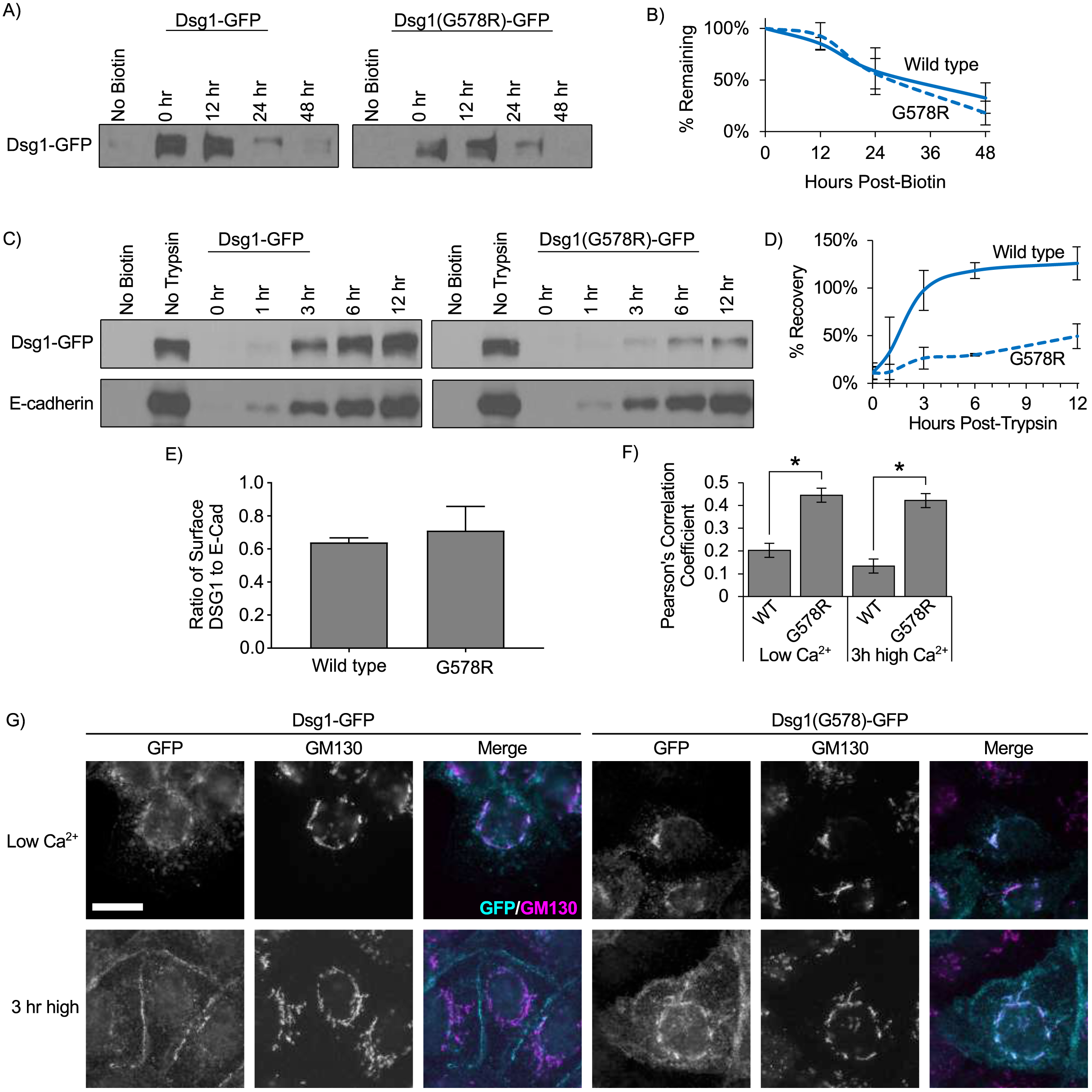
SAM-causing DSG1 mutation delays trafficking to the plasma membrane. A) Pulse chase experiments were preformed to determine the rate of turnover of Dsg1 from the plasma membrane in A431 cells expressing murine Dsg1-GFP. Cell surface proteins were biotinylated at t=0, washed and incubated at 37 degrees for various amounts of time. Cell lysates were collected after the indicated times. Biotinylated proteins were captured using streptavidin beads and processed for western blot analysis B) Quantification using densitometry revealed no significant differences in the rate of turnover of Dsg1 versus Dsg1(G578R) C) Dsg1(G578R) is trafficked to the plasma membrane substantially more slowly than wild-type. Cell surface proteins were cleaved using trypsin at t=0. Trypsin was removed and cells were then incubated for the indicated times. The amount of newly delivered surface Dsg1 was assayed via biotin labeling followed by capture using streptavidin beads and subsequent western blot analysis. D) Quantification using densitometry indicates Dsg1(G578R) recovers more slowly than WT-Dsg1. E) Densitometry analysis of the Dsg1 no-trypsin condition in panel C as a ratio to densitometry analysis of the E-cadherin no-trypsin condition in panel C reveals comparable surface levels of WT Dsg1 and Dsg1(G578R). F) Cells were cultured in low calcium medium to cause cadherin removal from cell-cell borders and accumulation in intracellular compartments (Panel E, Low Ca2+). Some cells were then switched back to normal calcium to allow for junction assembly (Panel E, 3 hr high). Dsg1(G578R) displays increased colocalization with the Golgi apparatus protein GM130 under both conditions. Scale bar = 20μm. G) Quantification of colocalization of Dsg1 and GM130. *p<0.001

### Disease causing mutation abrogates lipid raft association of Dsg1

Based on our findings that the TMD of the desmogleins is critical for lipid raft association, we hypothesized that the G-to-R mutation observed in SAM patients would prevent Dsg1 from partitioning to lipid rafts. Calculations based on parameters which predict free energy of raft association from TMD sequences [47] revealed that introduction of the SAM-causing G-to-R mutation into the DSG1 TMD dramatically alters the energetics of raft association (Table 1). Consistent with these calculations, sucrose gradient fractionations revealed that Dsg1(G578R) was virtually absent from lipid raft (DRM) fractions (Figure 7A and 7B). Interestingly, we observed a notable reduction in plakoglobin association with DRM in cell lines expressing Dsg1(G578R), suggesting that the DSG1 mutant also recruited plakoglobin out of raft domains. To further test the ability of WT Dsg1 and Dsg1(G578R) to associate with lipid rafts, these proteins were transiently expressed in rat basophilic leukemia cells and giant plasma membrane vesicles (GPMV) were chemically isolated [56, 57]. Non-raft plasma membrane domains were labelled with F-DiO, a dialkylcarbocyanine dye. WT Dsg1 efficiently partitioned into areas of plasma membrane vesicles lacking F-DiO, indicating partitioning to the liquid ordered, raft domain (Figure 7C and 7D). In contrast, Dsg1(G578R) was almost entirely co-segregated with F-DiO and excluded from the liquid ordered plasma membrane domain, indicating minimal raft affinity. Together, these findings reveal that the G-to-R TMD mutation reduces cell surface DSG1 association with lipid rafts.

**Figure 7:**
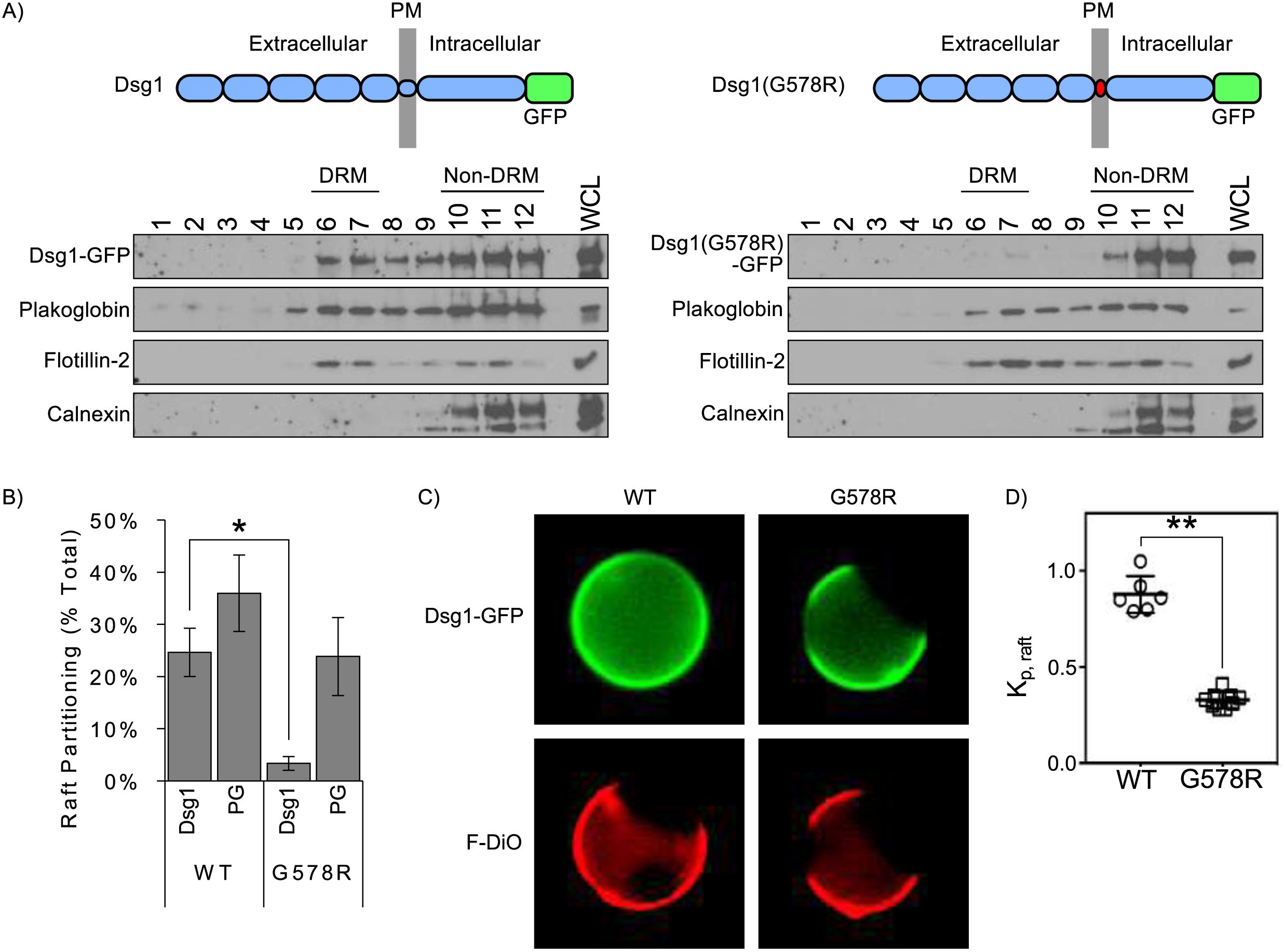
SAM-causing DSG1 mutation abolishes lipid raft association. A) Sucrose gradient fractionation and western blot analysis of A431 cell lines stably expressing WT and mutant Dsg1. B) Quantification of results in Panel A indicates SAM-causing mutation abolishes Dsg1 partitioning to DRM (lipid raft fractions). C) Representative images of giant plasma membrane vesicles isolated from rat basophilic leukemia cells expressing GFP-tagged WT Dsg1 or Dsg1(G578R). Unsaturated lipid marker FAST-DiO (F-DiO) to visualize the non-raft phase. D) Normalized line scans of Dsg1 fluorescence intensity were measured through peaks corresponding to Dsg1 intensity in raft and nonraft membrane, respectively. Background-subtracted ratios of these two intensities yield raft partition coefficients, K_p,raft_. Data are shown as mean ± SEM from three independent experiments. *p<0.05, **p<0.001

### The lipid bilayer within desmosomes is thicker than non-desmosomal membranes

The results above illustrate a critical role for the desmoglein TMD in the association of this family of cadherins with lipid raft membrane microdomains and for its crucial role in epidermal homeostasis. To understand how the physiochemical properties of the desmoglein TMD confer raft and desmosome targeting, structural models of the TMDs of wild type Dsg1, the Dsg1(G578R) SAM mutant, and E-cadherin were generated by the Robetta structure prediction server [58]. The modeling predicts that the SAM-causing G-to-R mutation interrupts the Dsg1 TMD helix and significantly shortens the run of helical hydrophobic residues (Table 1 and Figure 8A), potentially deforming the lipid bilayer as phospholipids position to maintain energetically favorable interactions between hydrophilic and hydrophobic amino-acid residues [59, 60]. These findings are consistent with the notion that the SAM causing mutant disrupts lipid raft association by shortening the DSG1 TMD, thereby increasing the energy cost of entering the thicker lipid bilayer characteristic of lipid raft domains [47].

**Figure 8:**
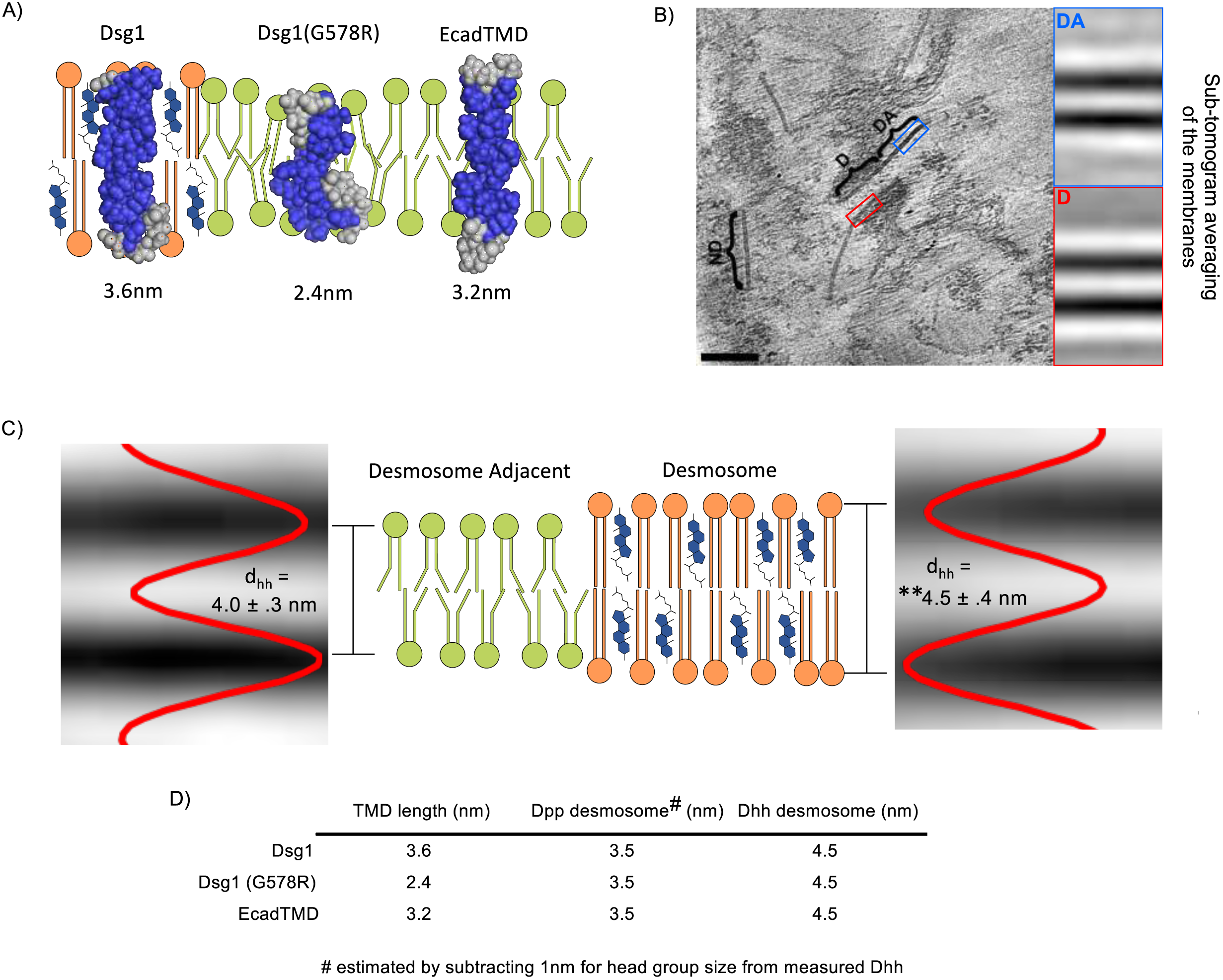
The desmosome bilayer is thicker than adjacent bilayers. A) Structural models of the DSG1 WT, DSG1 SAM mutant and E-cadherin TMDs acquired using the Robetta prediction server and depicted in schematized lipid bilayers. The length of each TMD is shown in nm and is based on TMD amino-acid number. B) Representative slice from a cryo-electron tomogram showing a desmosome (D) with characteristic intracellular plaque attached to intermediate filaments. Directly adjacent to the desmosome, membrane remnants can be seen (DA). Other non-desmosomal (ND) regions of the plasma membrane embedded in a thin layer of ice are also visible. Insets are projections of the average of all sub-volumes from the most significant class. Scale bar = 100nm. C) Schematic showing the thickness of the desmosome bilayer compared to desmosome adjacent bilayers. The lipid bilayer within desmosomal membranes is thicker (4.5 ± 0.4 nm) when compared to membranes adjacent to desmosomes (4.0 ± 0.3 nm) or from non-desmosomal membranes (4.0±0.3 nm) **p<.001. Intensity plots are shown superimposed to sub-tomogram average projections for desmosome and desmosome adjacent membranes. D) Summary table depicting the TMD lengths and the measured phospho-head group to head group distance (D_hh_) as shown in Panel C. Also shown is the estimated distance between phosphate residues (D_pp_) which corresponds to the hydrophobic interior of the bilayer. This hydrophobic region of the bilayer was estimated by subtracting the predicted polar head group size (1nm) [62] from the measured D_hh_ shown in Panel C.

Experiments in model membranes suggest that the high cholesterol content [59, 61] and long, saturated acyl chains [62, 63] present in lipid raft domains contribute to significant thickening of raft phospholipid bilayers relative to non-raft regions of the membrane. A prediction derived from such observations, and from the experimentally demonstrated presence of desmosomal proteins in rafts, is that the lipid bilayer within desmosomes in cells or tissues would be thicker than non-desmosomal regions of the plasma membrane, thereby accommodating the lengthy desmoglein TMD. To test this possibility, cryo-electron tomography and sub-tomogram averaging was performed on mouse liver samples enriched in the plasma membrane fraction (Figure 8B). The thickness of lipid bilayers measured in the sub-tomogram averages within desmosomal and non-desmosomal regions of the plasma membrane was then determined. This analysis revealed that desmosomal bilayers were 10% thicker (4.5±0.4 nm) than regions immediately adjacent to the desmosome (4.0±0.3 nm, p = 2.2E-122) or at arbitrary regions of membrane visible within the tomograms (4.0±0.3 nm, p = 6.4E-104) (Figure 8C). Together, these findings suggest that desmosomes represent a highly specialized plasma membrane domain that is characterized by lipid raft associated proteins and a thickened phospholipid bilayer characteristic of lipid raft-like model membranes.

## Discussion

Lipid rafts have emerged as important membrane microdomains that regulate membrane organization, endocytosis, and signaling [31, 33–36]. Desmosomal proteins have been shown to associate with lipid rafts in a variety of epithelial cell types [25–29], but the mechanisms and physiological relevance of this association are poorly understood. Here, we report that the transmembrane domains of the desmogleins are the key determinants for targeting these cadherins to lipid rafts. A mutation within the DSG1 TMD that shortens this domain abrogates both lipid raft partitioning and desmosome association, and leads to the human skin disease SAM syndrome. Cryo-electron tomography reveals that the lipid bilayer within the desmosome is markedly thicker than the adjacent lipid bilayer, thereby favoring incorporation of the longer desmoglein TMDs into this plasma membrane domain. Collectively, our results suggest that desmosomes are a specialized mesoscale lipid raft-like membrane domain.

Essential functions for desmogleins have been exposed by human diseases in which desmogleins are targeted by autoantibodies, infectious agents, or genetic mutation [3, 22]. DSG1 is the primary desmoglein expressed in the outermost layers of the epidermis, and DSG1 loss of function mutations lead to at least two different types of epidermal disorders. Haploinsufficiency of DSG1 causes palmoplantar keratoderma [64], whereas complete loss of DSG1 leads to SAM syndrome [24, 52, 53]. Most individuals afflicted with SAM syndrome succumb to chronic infection in early childhood [24]. Here, we report two separate instances of SAM syndrome, one inherited and one sporadic, caused by a glycine to arginine substitution (G562) within the hydrophobic transmembrane domain (Figure 4). Arginine residues play an important role in terminating TMDs and establishing TMD orientation within the lipid bilayer [65, 66], consistent with molecular modeling indicating that the disease-causing glycine to arginine substitution shortens the DSG1 TMD (Fig 8).

Our data indicate that shortening of the DSG1 TMD by insertion of an arginine residue disrupts DSG1 function in SAM syndrome patients by preventing lipid raft association. TMD length correlates positively with raft association [47], and our structural predictions and molecular modeling predict that desmoglein TMDs confer raft association (Table 1 and Figure 8A). This notion is consistent with our findings using both classical DRM fractionation experiments (Figure 3 and 4) and direct observations of partitioning of cell surface Dsg1 into liquid ordered plasma membrane domains (Figure 7). Surprisingly, palmitoylation does not appear to be required for desmoglein raft association (Figure 1 and reference 45), although it does impact desmoglein dynamics at the plasma membrane [45]. Recently, the desmosomal component plakophilin also was shown to be palmitoylated [40], further demonstrating an important role for this reversible posttranslational modification in regulating desmosome assembly dynamics. Further studies will be needed to assess how palmitoylation is utilized in combination with other physiochemical properties of the desmoglein TMD to modulate the trafficking and adhesive properties of these unique cadherins.

A prediction based on our finding that the desmoglein TMD is responsible for partitioning to lipid rafts is that the lipid bilayer within desmosomes should be thicker than surrounding non-desmosomal membrane. Indeed, cryo-electron tomography revealed that the lipid bilayer within the desmosome is substantially thicker than non-desmosomal regions of the plasma membrane (Figure 8B-8D). These observations indicate that it would be energetically costly for the DSG1 G-to-R SAM mutant to enter the thicker bilayer present in desmosomes due to hydrophobic mismatch between phospholipids and Dsg1 TMD amino-acid residues [60, 67]. Therefore, it is likely that shortening of the TMD in the SAM mutant and failure to enter the thicker lipid bilayer domain of the desmosome represents a central pathophysiological mechanism of this disease-causing mutation. Indeed, we observed that the DSG1 G-to-R mutant is deficient in entering desmosomes both in patient epidermis (Figure 4) and when expressed in cultured epithelial cell lines (Figure 5). We also find that Dsg3 and Dsg1 polypeptides harboring the shorter E-cad TMD are unable to associate with lipid rafts and behave similarly to full length E-cadherin (Figure 2, Figure 3). The predicted E-cadherin TMD is 21 amino acids, compared to the 24 amino acid TMD of desmogleins (Table 1). Although these chimeras do not effectively enter lipid rafts as assessed by DRM fractionation assays, we do find that these Dsg(EcadTMD) chimeras can associate with desmosomes as assessed by SIM (not shown). It is likely that for these chimeras, protein-protein interactions mediated by the desmoglein cytoplasmic and extracellular domains can partially overcome the energy cost of incorporating into the thicker bilayers present in the desmosome. In addition, mismatch of TMD length and hydrophobic thickness of the bilayer can be accommodated by changes in TMD tilt within the membrane and by local bilayer deformation [59, 60]. In contrast, the predicted 16 amino acid TMD of the Dsg1(G578R) mutant is significantly shorter than the TMD of both desmogleins and E-cadherin, and therefore its entry into desmosomal membranes is apparently energetically prohibitive.

Together, our observations support a model in which adherens junctions and desmosomes assemble into distinct plasma membrane microdomains based not only on protein interactions, but also due to the biophysical nature of the epithelial plasma membrane and the TMD characteristics of different cadherin subfamilies (Figure 9A and 9B). Interestingly, early studies of desmosomal composition found that these junctions are enriched in sphingolipids and cholesterol, key components of what are now referred to as lipid rafts [68, 69]. In addition, most of the major desmosomal proteins are palmitoylated, including the desmosomal cadherins and plakophilins, whereas adherens junction components lack this modification [40, 45]. Given the key role for palmitoylation in lipid raft association, these findings further suggest that affinities for different lipid domains of the plasma membrane are central features that distinguish adherens junction and desmosomal proteins. Further studies will be needed to determine the precise structural and functional characteristics of different cadherin TMDs and how they selectively dictate lipid raft association. In addition, it will be important to discern how TMD characteristics are used in conjunction with lipid modifications such as palmitoylation to sort desmosome and adherens junction components into distinct plasma membrane domains with unique morphologies and functions. These features appear to be of fundamental importance for skin physiology, as our findings reveal that a mutation altering the structure of the desmoglein transmembrane domain is a novel pathomechanism of a desmosomal disease. This work also raises the possibility that other human disorders may result from alterations in lipid raft association or raft homeostasis. Indeed, loss of lipid raft targeting may be an under-appreciated pathomechanism in human diseases which were previously conceived as generalized protein trafficking defects.

**Figure 9:**
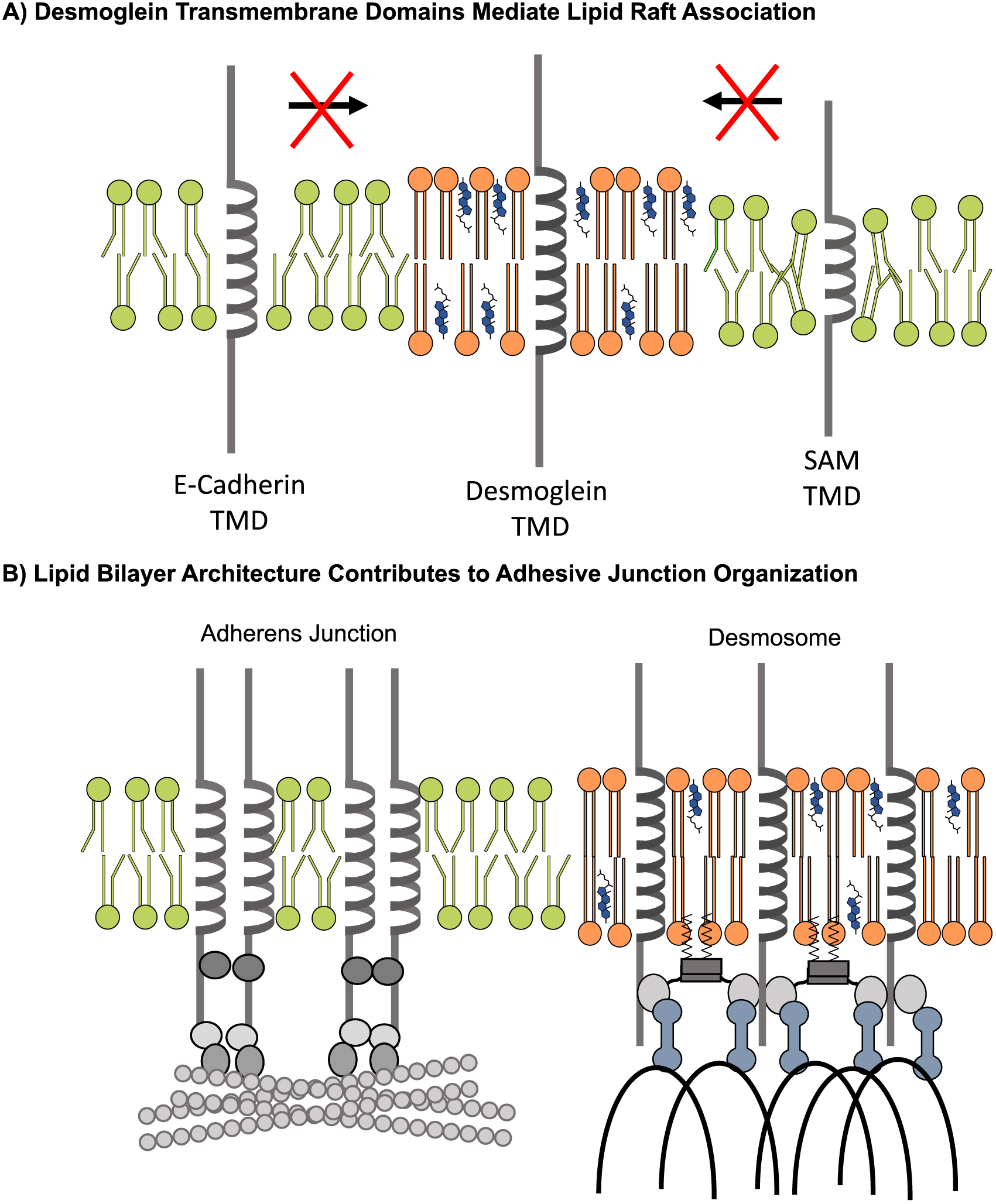
Model. A) I. The extended lenth of the desmoglein transmembrane domain aids in the recruitment and partitioning to sphingolipid- and cholesterol-enriches vesicles budding off of the Golgi network. II. Once desmoglein is at the plasma membrane, the transmembrane domain length regulates association and incorporation to the larger desmosomal complex B) The extended desmoglein transmembrane domain facilitates lipid raft association. In contrast, the entry of E-cadherin and the Dsg1 SAM mutant into lipid rafts is unfavorable due to hydrophobic mismatch between the cadherin TMD and the phospholipid headgroups of the lipid bilayer. C) Desmosomal proteins enter lipid raft domains through TMD affinities for raft-like membrane domains and palmitoylation of desmosomal cadherins and plaque proteins. In contrast, adherens junction components lack these raft-targeting features, resulting in exclusion of adherens junction components from lipid raft membrane domains. Thus, the biophysical properties of the bilayer associated with the desmosome promote spatial segregation of adherens junctions and desmosomes

## Acknowledgements

The authors would like to thank Dr. Kathleen Green and members of the Kowalczyk lab for comments and insights during the preparation of this manuscript. We would also like to thank Joseph Lorent for help generating the transmembrane domain models. This work was supported by grants (R01AR048266, R01AR048266-13S1, and R01AR050501 to A.P.K.), (LOEWE Dynamem to A.S.F) and fellowships (F31AR066476 and T32GM008367 to J.D.L.) from the National Institutes of Health, and by the Practical Research Project for Rare/Intractable Diseases (16ek0109067h0003 to Y.M. and 16ek0109151h0002 to A.K.) from the Japan Agency for Medical Research and Development. Additional support was provided by core facilities at Emory University, including the Integrated Cellular Imaging core, the Emory Flow Cytometry Core, and the Cloning Division within Emory Integrated Genomics Core.

The authors declare no competing financial interests.

## Author Contributions

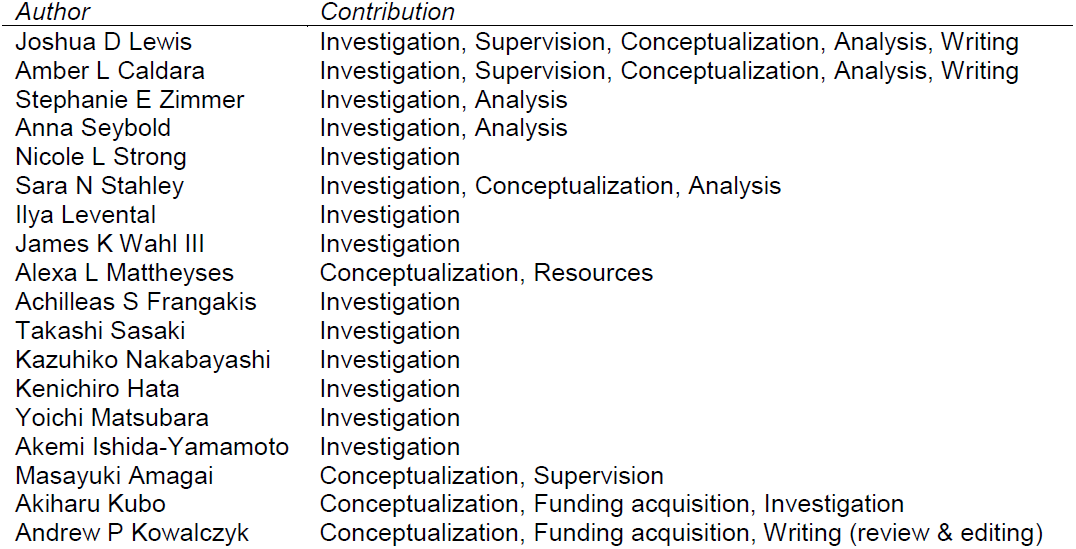

## Materials and methods

### Subjects

All affected and healthy family members or their legal guardians provided written and informed consent in accordance with the guidelines of the Institutional Review Board of Keio University and Emory University School of Medicine in adherence to the Helsinki guidelines. The investigators were not blinded to the allocation during experiments and outcome assessment.

### Mutation analysis

Whole-exome sequencing was performed using genomic DNA isolated from the probands (II-2 and III-2) and their parents (I-1, I-2 and II-1). Whole exome sequencing libraries were constructed using SureSelect Human All Exon V5 (Agilent) and sequenced by HiSeq2500 (Illumina). Sequencing reads were mapped to a human reference genome sequence (hs37d5) by BWA software (0.7.12-r1039). The mapped reads were realigned and variation sites were detected by GATK-3.30 software. The detected variation sites were annotated by SnpEff/SnpSift 4.1d software. Since the phenotype appeared in the proband II-2 (delivered from healthy parents) and transmitted to the proband III-2 (Figure 4D), we searched for genetic variations that de novo mutated in the proband II-2 and transmitted to the proband III-2. Only one variation was identified to fulfill the criteria, which was c.1684G>A (p.G562R) of *DSG1*, coding for the desmosomal cadherin desmoglein 1. Sanger sequencing confirmed the mutation was identified in the probands but not from other healthy family members (Figure 4D, 4E). The mutation had not been identified in cohort studies [70–73]. The whole exome sequencing of the probands II-2 and III-2 revealed no other variations in the exons and exon-intron boundaries of *DSG1*.

### Immunohistochemistry and electron microscopy of patient samples

Biopsies were embedded in optimum cutting temperature (OCT) solution and stored at −80°C. Prior to immunostaining, 5 μm cryosections were prepared on glass microscope slides. Primary and secondary antibodies are described below. Sections were sealed using mounting medium (ProLong Gold by ThermoFisher Scientific) and a coverslip. For electron microscopic studies, the biopsied sample was fixed in an ice-cold 2% glutaraldehyde/60 mM Hepes (pH 7.4) buffer followed by fixation with 1% osmium tetroxide, staining with 1% uranyl acetate, and embedding in Epon812. Ultrathin sections were stained with 1.5% uranyl acetate and Reynolds lead citrate and examined with an electron microscope (JEM-1010, JEOL) at the accelerating voltage of 80 kV.

### Construction of mutants

Constructs were cloned using PCR and mutagenesis by the Cloning Division within Emory Integrated Genomics Core or purchased through Cyagen vectorbuilder services.

### Structural Predictions

Sequences for transmembrane domains were analyzed using the Robetta structure prediction server [58]

### Cell line generation, culture, and reagents

A431 cells were cultured in DMEM (Corning 10- 013-CV) with 10% fetal bovine serum (Hyclone SH30071.03) and 1% penicillin/streptamycin (Corning 30-004-CI). Cells were stably infected with lentiviruses expressing the various murine desmoglein constructs. 5 μg/mL blasticidin was used to select for infected cells. No clonal isolation was performed. Cell lines expressing wild type and mutant DSG1-GFP were subjected to fluorescence activated cell sorting in order to obtain populations with roughly equal DSG1-GFP expression levels. For experiments utilizing a calcium switch, low calcium medium was prepared as described previously [74]: no calcium DMEM (Gibco/Molecular Probes 21068028), 10% fetal bovine serum, calcium chelating BT Chelex 100 resin (Biorad 143-2832), and 1% penicillin/streptamycin.

### Immunofluorescence

A431 cells were cultured to ~70% confluence on glass coverslips. In experiments in which pre-extraction is explicitly used, cells were treated with PBS+ containing 0.2% Triton X-100 and 300 mM sucrose on ice for 1 min prior to fixation. Cells were fixed in 3.7% paraformaldehyde in PBS+ on ice for 10 min. Cells were permeabilized in PBS+ containing 0.1% Triton X-100 and 3% bovine serum albumin for 10 min. Non-specific antibody binding was prevented with a blocking step in PBS+ containing 3% bovine serum albumin and 0.05% Triton X-100. Primary and secondary antibodies (listed below) were diluted into blocking solution. For rinse buffer, we used PBS+ containing 0.2% bovine serum albumin and 0.05% Triton X-100. Cells were mounted to glass microscope slides using prolong gold mounting medium (described above).

### Antibodies

Mouse anti-DSG3 AK15 was described previously [75]. Rabbit anti-calnexin (Enzo Life Sciences ADI-SPA-860). Mouse anti-desmoplakin1/2 (Fitzgerald 10R-D108AX). Rabbit anti-desmoplakin NW6 was a kind gift from Dr. Kathleen Green (Northwestern University). Mouse anti-plakoglobin (gamma catenin) (BD TransLabs 610253). Mouse anti-E-cadherin (BD Biosciences 610252). Mouse anti-flotillin 1 (BD 610820). Mouse anti-flotillin 2 (BD 610383). Rabbit anti-Green Fluorescent Protein Life A11122). Rabbit anti-FLAG (Bethyl A190-102A). Secondary antibodies conjugated to Alexa Fluors were purchased from Invitrogen. Horseradish peroxidase-conjugated secondary antibodies were purchased from Biorad.

### Image acquisition and processing

Widefield fluorescence microscopy was performed using a DMRXA2 microscope (Leica, Wetzler, Germany) equipped with a 100X/1.40 NA oil immersion objective and narrow band pass filters. Images were acquired with an ORCA digital camera (Hamamatsu Photonics, Bridgewater, NJ) and processed using Fiji ImageJ. Super-resolution microscopy was performed using a Nikon N-SIM system on an Eclipse Ti-E microscope system equipped with a 100X/1.49 NA oil immersion objective, 488- and 561-nm solid-state lasers in 3D structured illumination microscopy mode. Images were captured using an EM charge-coupled device camera (DU-897, Andor Technology) and reconstructed using NIS-Elements software with the N-SIM module (version 3.22, Nikon). All microscopy was performed at room temperature and imaging results are representative for at least two independent experiments containing at least 10 cells each.

### Desmosome targeting analysis using SIM

To quantify desmosome targeting in cultured cells, Dsg1.GFP fluorescence was measured within regions of interest (ROI) drawn around desmoplakin railroad track staining at cell-cell borders. This Dsg1.GFP fluorescence intensity was compared to adjacent ROI at regions of cell borders lacking desmosomes. For both wild type and mutant Dsg1, targeting to desmosomes was measured as a fold-enrichment of Dsg1.GFP fluorescence in desmosomes compared to non-desmosomal regions. For SAM patient and control tissue, desmosomal ROIs were defined using desmoplakin railroad tracks and DSG1 fluorescence was measured therein.

### Triton solubility/insolubility

A431 cells were cultured until confluent in 6 well tissue culture plates. Cells were washed twice with ice cold phosphate buffered saline. The triton soluble pool was isolated by incubating cells with triton buffer (1% Triton X-100, 10 mM Tris, pH 7.5, 140 mM NaCl, 5 mM EDTA, 2 mM EGTA, with protease inhibitor) for 10 min on ice. Lysate was then centrifuged at 16,000 x *g* for 10 min at 4°C to pellet triton insoluble fraction. Triton-soluble supernatant was collected and mixed 1:1 with 2x laemmli sample buffer containing 5% B-mercaptoethanol. The triton-insoluble pellet was resuspended in 2X laemmli sample buffer (Biorad 161-0737) sample buffer containing 5% β-mercaptoethanol. All samples were heated to 95°C for 10 minutes, vortexed for 30s half way through, prior to being run on a gel for western blotting.

### Isolation of detergent resistant membranes

Detergent resistant membranes were isolated as described previously [46]. Briefly, cells were cultured in 25 cm^2^ flasks (two per gradient) and washed with PBS+. Cells were collected by scraping in TNE buffer supplemented with protease inhibitors (Roche) and pelleted by centrifugation at 0.4 x *g* at 4°C for 5 min (5415R, Eppendorf). Cells were re-suspended in TNE buffer and homogenized using a 25-guage needle. TNE buffer containing Triton X-100 was added (final concentration of 1%) and cells were incubated on ice for 30 min. 400 μL of detergent extract was mixed with 800 μL of 56% sucrose in TNE and placed at the bottom of a centrifuge tube. 1.9 mL volumes of 35% and 5% sucrose were layered on top of the sample. Following an 18 hour centrifugation at 4°C (44,000 rpm, SW55 rotor, Beckman Optima LE-80 K Ultracentrifuge), 420 μL fractions (1–11, remaining volume combined to make up fraction 12) were removed from top to bottom of the gradient and stored at −20°C until processed for western blot analysis. Flotillin-1 and Flotillin-2 were used as raft markers while calnexin was used as a non-raft marker. Unless otherwise stated, all films shown are representative for at least three independent experiments.

### Giant plasma membrane vesicle (GPMV) isolation and partitioning measurements

GPMVs were isolated and imaged as described [76, 77]. Before GPMV isolation, cell membranes were stained with 5 μg/mL of FAST-DiO (Invitrogen), a fluorescent lipidic dye that strongly partitions to disordered phases because of double bonds in its fatty anchors [78].

### Biotin labeling in pulse-chase experiments

For Dsg1 cleavage and recovery experiments, cells were grown to confluence in 35 mm cell culture plates (Corning 430165). Cells were trypsinized using TrypLE (Gibco 12605-010) for ~8 min and suspended. After the indicated refractory period, surface proteins were biotinylated. For experiments monitoring protein turnover from the plasma membrane, surface proteins were biotinylated before the indicated period. Biotinylation was achieved using PBS+ containing 0.5 mg/ml EX-Link sulfo NHS SS Biotin (Thermo Scientific 21331) for 30 min at 37°C. Unbound biotin was quenched in PBS+ containing 50 mM NH_4_Cl for 1 min. Cells were lysed in RIPA (PBS+ containing 1% Triton X-100, 0.1% sodium dodecyl sulfate, 0.1% sodium deoxycholate, 10 mM Tris-HCl, 140 mM NaCl, 1 mM EDTA, 0.5 mM EGTA, and protease inhibitor cocktail (Roche 11836170001)), scraped to transfer from culture plate to an Eppendorf tube, and incubated for 10 min on ice. Lysate was cleared via centrifugation at 16,000 x *g* at 4°C for 10 min. Biotinylated protein was captured on streptavidin-coated beads (manufacturer) during overnight incubation at 4°C. Beads were collected via centrifugation at 2,500 x *g* at 4°C for 1 min. Protein was released from beads using Laemmli buffer containing 5% β-mercaptoethanol.

### Mass-tagging of palmitoylated proteins

For mass-tag labeling, we followed the procedure described by [79]. Lysates from A431 cells expressing the indicated constructs were prepared in TEA buffer (50 mM triethanolamine; pH7.3, 150 mM NaCl, and 5 mM EDTA) containing 4% SDS. 200 μg of total cellular protein was treated with a final concentration of 10 mM neutralized TCEP for 30 min with end over end rotation. NEM was added to a final concentration of 25 mM and rocking continued for 2 hours. NEM was removed by 3 rounds of chloroform/methanol/H_2_O precipitation. The final pellet was resuspended in TEA buffer containing 0.2% Triton X-100. Samples are treated with 0.75 M NH_2_OH (+HA) or without hydroxylamine (-HA) and incubated at room temperature for 1 hour. Excess hydroxylamine was removed with one round of chloroform/methanol/H_2_O precipitation and the pellet was resuspended in TEA buffer containing 0.2% Triton X-100 supplemented with 1 mM mPEG-Mal (10 kDa; Sigma). Samples were incubated with rocking for 2 hours and reactions were terminated by 1 round of chloroform/methanol/H_2_O precipitation. The final pellet was suspended in 1x Laemmli sample buffer and resolved by SDS-PAGE.

### Isolation, freezing, and imaging of desmosomes

To isolate desmosomes from mouse liver, a method based on the protocol of Tsukita and Tsukita (1989) was used, in which a desmosomal fraction was obtained by sucrose density gradient centrifugation, followed by NP-40 detergent treatment. The fraction should contain only bile canaliculi derived plasma-membranes, as the homogenization and centrifugation steps are designed to free the preparation from contaminating cell fragments and nuclear membranes due to their higher densities [80–82]. The desmosomal fraction was immediately plunge-frozen on holey carbon grids, which were subsequently inserted into the column of a FEI Titan Krios at liquid nitrogen temperatures. Tilt series of the sample (+60 to −60°) were recorded and subsequently reconstructed into 3D tomograms.

### Membrane thickness measurements

For the thickness of the desmosomal membranes, 1768 selected positions (derived from 3 desmosomes in 2 tomograms) with visible cadherins were selected. As a comparison (control), 668 randomly selected positions at membranes adjacent to desmosomes (derived from 3 membranes in 3 tomograms) and 515 randomly selected positions from arbitrary membranes in the tomograms (derived from 2 membranes in 1 tomogram) were selected. Each position was cross-correlated with multiple references of a simplified membrane model of the two leaflets (dark lines representing phospholipid head groups and are included in the measurements) with varying bilayer distance (3.08, 3.52, 3.96, 4.4, 4.84 and 5.28 nm spacing) using sub-tomogram averaging routines with limited rotational freedom (±30° in 5° steps for all three Euler-angles) after rough pre-alignment using the overall membrane orientation. The reference with the highest cross-correlation score then provides the bilayer spacing of each single sub-volume

### Statistics

Error bars represent standard error of the mean. Significance was determined using a student’s t-test (two tailed, heteroscedastic) and p-values have been indicated. Statistical analysis of immunofluorescence results was conducted on at least two independent experiments with ten images per condition per replicate. Statistical analysis of western blotting was conducted on results from three independent experiments.

